# The roles of RNA editing in cancer immunity through interacting with interferon

**DOI:** 10.1101/2023.08.06.552142

**Authors:** Sijia Wu, Xinyu Qin, Zhennan Lu, Jianguo Wen, Mengyuan Yang, Pora Kim, Xiaobo Zhou, Liyu Huang

**Author notes:** Correspondence (X.Z.) and (L.H.).

## Abstract

The interferon-activated tumor innate immunity can be primed by specific double-stranded RNA (dsRNA) sensors upon stimulation. A-to-I RNA editing in the dsRNA regions can have a potential function to regulate interferon-related cancer immunity. A systematical analysis of both the editing enzyme and specific enriched editing region in patients, tissues, and cell lines is performed to reveal the underlying mechanisms. We then validate the preferred editing of dsRNA regions, identify the hyper-editing in severe tumors, and discover the negative effect of editing on cancer immunity. Specifically, RNA editing acts as an inhibitor of *PKR*- and *MDA5*-related interferon pathways through the regulations of miRNAs and RNA-binding proteins and the deactivation of dsRNA sensors. With the alteration of interferons, subsequently, RNA editing represses the infiltration of CD8 and CD4 T cells and reduces the sensitivities of cancer drugs, such as cisplatin. These analyses on A-to-I RNA editing can improve the knowledge of tumorigenesis, immunology, and cancer-targeted immunotherapy.

**Highlights:**

1. The preferred dsRNA region for RNA editing is validated.
2. Upregulation of RNA editing in severe tumors is discovered.
3. RNA editing inhibits PKR- and MDA5-related cancer immunity.
4. RNA editing represses the infiltration of CD8 and CD4 T cells.
5. RNA editing reduces the sensitivities of cancer drugs.

## Introduction

Therapies that promote antitumor immunity revolutionize cancer treatment (1). Most of these immunomodulatory approaches focus on enhancing T-cell responses (2). Interferons are essential for the priming of tumor-specific T cells, thereby contributing to tumor control and immunoediting (3,4). However, chronic and persistent stimulation of the interferons can induce resistance to various anti-tumor therapies (5). Understanding the mechanism of interferon pathways is crucial for cancer therapies.

The families of double-stranded RNA (dsRNA) sensors stand out due to their close associations with interferon (6,7). For example, RIG-I like receptors, including melanoma differentiation-associated gene 5 (*MDA5*, also known as *IFIH1*), recognize dsRNAs to activate the common downstream adaptor molecule of mitochondrial antiviral signaling protein (*MAVS*), leading to the transcriptional upregulation of type I interferons (8). Protein kinase R (*PKR*, also known as *EIF2AK2*) consists of dsRNA-binding domains and is stimulated by interferon (9). These descriptions display the roles of dsRNA sensors in interferon-related biological functions. The dsRNA regions where these proteins bind have clusters of A-to-I RNA editing events (10). Then A-to-I RNA editing has great potential to participate in the regulations of interferon pathways, thus affecting the effectiveness of cancer immunotherapies (11,12).

To perform a comprehensive study about the roles of A-to-I RNA editing in interferon pathways, 11,052 patients across 33 cancer types from The Cancer Genome Atlas (TCGA), kidney normal and tumor tissues from GEO (13), *ADAR* knockdown cell lines from ENCORE project are all involved in this study. These samples are first analyzed for the quantification of *ADAR* expressions, the definition of enriched editing regions, and the estimation of their editing levels. After that, the editing-related data is applied to validate the preferred dsRNA region for editing events and the tumor-specific properties of edited regions. The associations of A-to-I RNA editing with tumor and dsRNAs guide the study of editing roles in dsRNA-participated interferon pathways in cancers. To realize the goal, the research on both *ADAR* and specific enriched editing regions is conducted from their associations with interferon-stimulated genes, interferon response pathways, T-cell infiltration, and drug sensitivities. The four aspects are highly relevant to interferon biological functions. These analyses can reveal the underlying mechanisms of RNA editing during interferon pathways in cancers and provide potential novel editing-related cancer therapy method.

## Results

### 1. The preferred dsRNA regions for A-to-I RNA editing

Our pipeline identifies 17,689 enriched editing regions on 11,052 samples across 33 cancer types (Figure 1A). Of all these edited regions, more than 94% align to non-coding domains and Alu repeats. (Figure 1B-C). These regions belong to 7,772 different genes. Compared to the genes without clusters of RNA editing, these genes tend to be longer (Figure 1D). This phenotype still exists when considering the introns, exons, and 3’-UTRs. It means that non-coding domains and Alu repeats of long genes (introns, exons, and 3’-UTRs) are more likely to be edited.

**Figure 1.**
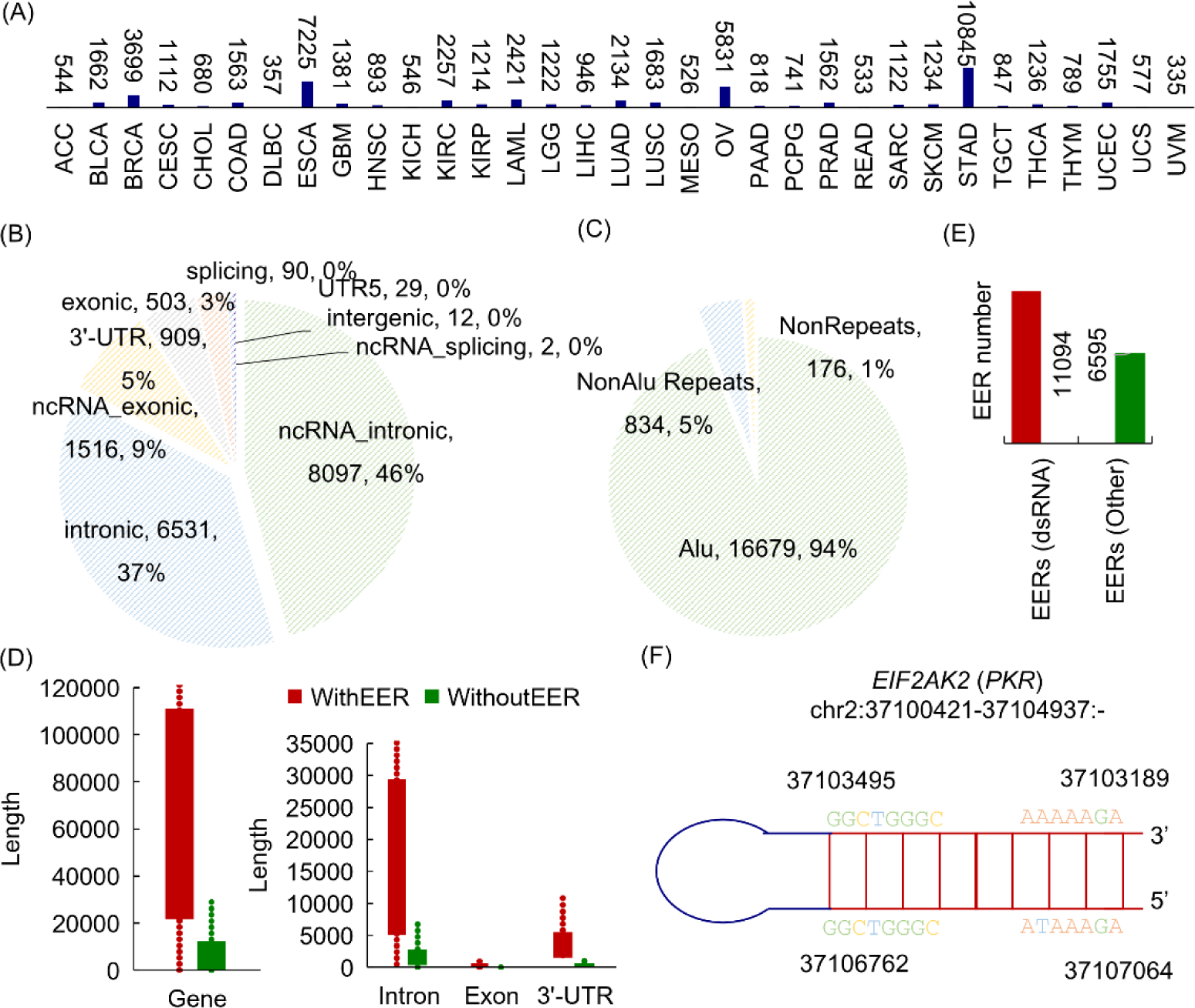
The preferred genomic distributions for A-to-I RNA editing. (A) The number of enriched editing regions identified in 33 cancer types. (B) The distributions of enriched editing regions in genomic locations. (C) The distributions of enriched editing regions in repeats. (D) The comparison of genes, introns, exons, and 3’-UTRs with enriched editing regions to that without enriched editing regions. The non-enriched editing genes were randomly selected to constitute a dataset that had an equal size with the dataset of enriched editing genes, similarly for introns, exons, and 3’-UTRs. (E) The distributions of enriched editing regions in dsRNA structures. (F) One edited region of *PKR* showing potential dsRNA structure with surrounding sequences. ACC, Adrenocortical Carcinoma; BLCA, Bladder Urothelial Carcinoma; BRCA, Breast Invasive Carcinoma; CESC, Cervical Squamous Cell Carcinoma and Endocervical Adenocarcinoma; CHOL, Cholangiocarcinoma; COAD, Colon Adenocarcinoma; DLBC, Lymphoid Neoplasm Diffuse Large B-Cell Lymphoma; ESCA, Esophageal Carcinoma; GBM, Glioblastoma Multiforme; HNSC, Head and Neck Squamous Cell Carcinoma; KICH, Kidney Chromophobe; KIRC, Kidney Renal Clear Cell Carcinoma; KIRP, Kidney Renal Papillary Cell Carcinoma; LAML, Acute Myeloid Leukemia; LGG, Brain Lower Grade Glioma; LIHC, Liver Hepatocellular Carcinoma; LUAD, Lung Adenocarcinoma; LUSC, Lung Squamous Cell Carcinoma; MESO, Mesothelioma; OV, Ovarian Serous Cystadenocarcinoma; PAAD, Pancreatic Adenocarcinoma; PCPG, Pheochromocytoma and Paraganglioma; PRAD, Prostate Adenocarcinoma; READ, Rectum Adenocarcinoma; SARC, Sarcoma; SKCM, Skin Cutaneous Melanoma; STAD, Stomach Adenocarcinoma; TGCT, Testicular Germ Cell Tumors; THCA, Thyroid Carcinoma; THYM, Thymoma; UCEC, Uterine Corpus Endometrial Carcinoma; UCS, Uterine Carcinosarcoma; UVM, Uveal Melanoma.

Most importantly, nearly 65% of enriched editing regions constitute dsRNA structures with their surrounding fragments (< 1e4 bp, Figure 1E-F). This number may increase when considering distant sequences. These self-derived dsRNAs are potential binding substates of cellular dsRNA sensors, such as RIG-I-like receptors (RLRs: *MDA5*) and protein kinase R (*PKR*) (14). The high tendency of A-to-I RNA editing in these dsRNA regions urges the study of RNA editing in dsRNA-sensing cancer immunity (11).

### 2. Upregulated RNA editing in severe tumor conditions

*ADAR* is the main enzyme responsible for the deamination of adenosine to inosine. Then the study of A-to-I RNA editing starts with differential *ADAR* expression analysis. For 15 cancer types with enough adjacent normal controls from the TCGA (> 10), all none-kidney tissues show increased *ADAR* expressions for tumors (*P* < 0.05, Figure 2A). The kidney tissues from the GEO database also have an overexpression of *ADAR* for tumors (*P* < 0.001, Figure 2B) (13). These abnormally up-regulated expressions suggest *ADAR* as a pathogenic biomarker of cancer.

**Figure 2.**
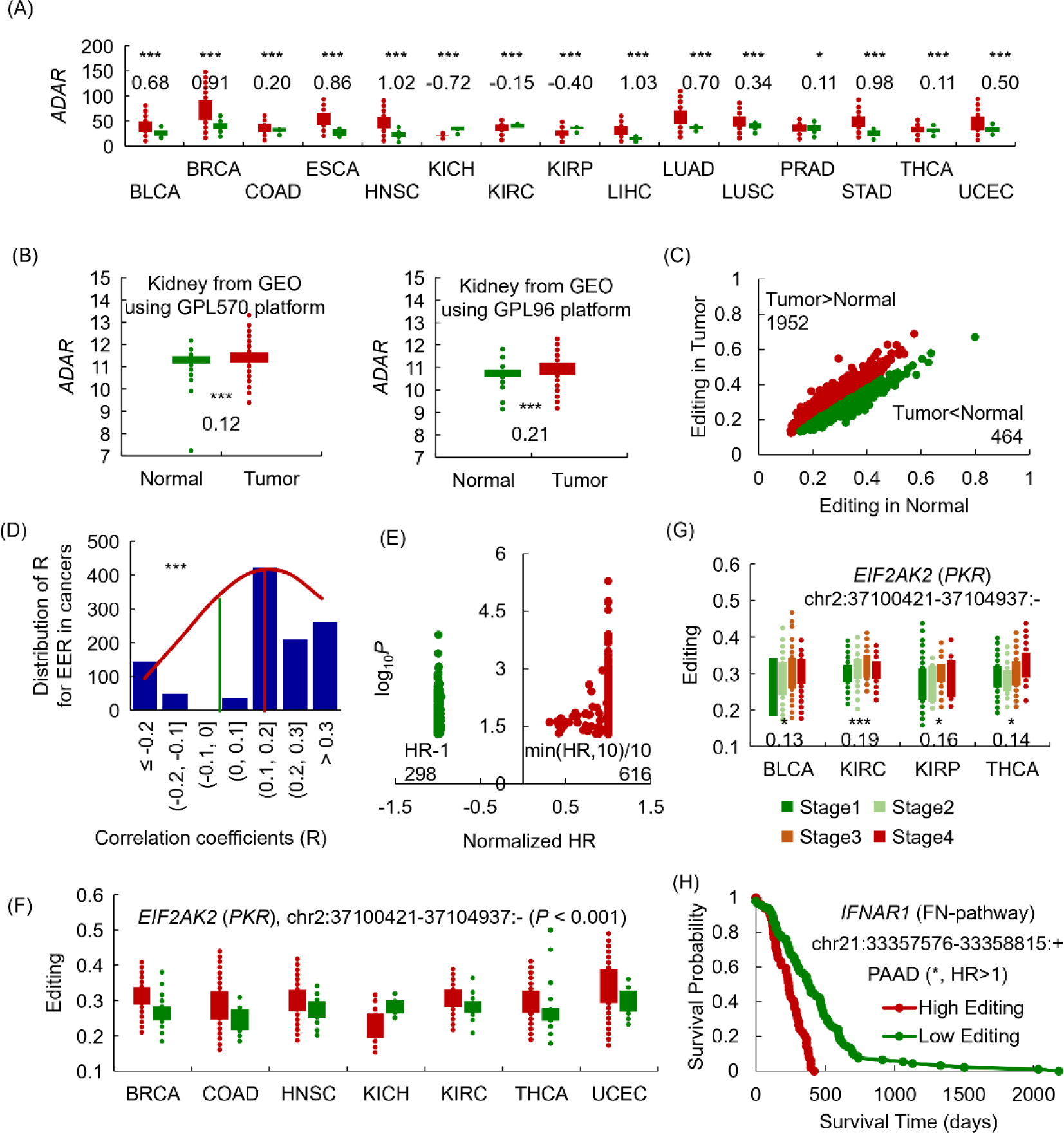
The changes of A-to-I RNA editing in tumors. (A) The comparison of *ADAR* expression between tumors (red) and controls (green) for 15 cancer types. The number in this panel is the log2FC value. (B) The comparison of *ADAR* expression between kidney tumors and controls from the GEO dataset. (C) The average editing levels of differential edited regions in tumor patients and normal controls. (D) The distribution of correlation coefficients for stage-associated edited regions. (E) The distribution of HR for survival-related edited regions. (F) The comparison of one enriched editing region of *PKR* gene between tumors (red) and controls (green). (G) The comparison of one enriched editing region of *PKR* gene among different stages. (H) The survival analysis result of one enriched editing region of *IFNAR1* gene in PAAD cancer type. *, P<0.05; **, P<0.01; ***, P<0.001; HR, hazard ratio.

The analyses on specific enriched editing region are also consistent with *ADAR* expression result. Specifically, more regions have statistically higher editing levels in tumors compared to controls (Figure 2C). The correlation coefficients between editing levels and pathological stages are significantly greater than zero (Figure 2D). There are beyond two-fold survival-related regions showing higher editing levels in poor survival groups (Figure 2E). These results introduce the increased overall editing level in tumors, severe cancer patients, and patients with high survival risks, such as the enriched editing regions of *EIF2AK2* (*PKR*) and *IFNAR1* gene (Figure 2F-H).

The abnormal upregulation of RNA editing reveals a strong relevance of RNA editing to cancer occurrence and progression. A close look at the genes with these tumor-specific enriched editing regions selects 70 genes associated with innate immunity (15), including the famous *PKR* (16) and *IFNAR1* genes (17) described above. It guides us to study the role of A-to-I RNA editing in cancer immunity, especially considering the preferred dsRNA regions for A-to- I RNA editing (Figure 1E) and the dsRNA-participated innate immunity (6).

### 3. The interplay between A-to-I RNA editing and immune genes

A comprehensive analysis of enriched editing regions and genes identifies 911 innate immune genes from InnateDB (15) potentially involving A-to-I RNA editing. Thus, many immune functions are affected, including the immunotherapy-targeted T-cell and PD-L1 pathways (Figure 3A). It describes the necessity of studying the editing roles in cancer immunity.

**Figure 3.**
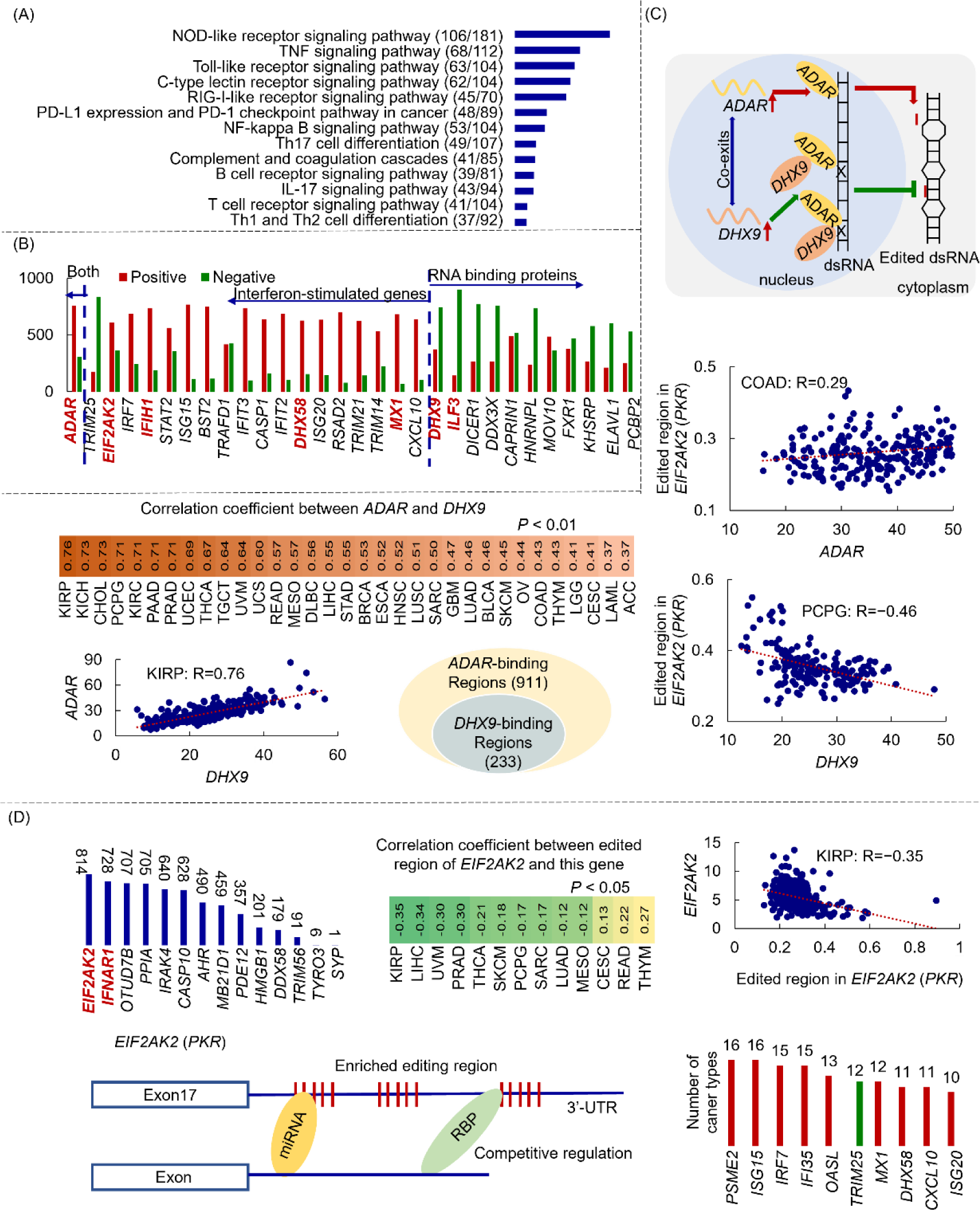
The analysis between edited regions and immune genes. (A) The biological functions of the editing-associated immune genes. (B) The number of edited regions positively (red) and negatively (green) associated with these interferon-stimulated genes or RNA-binding proteins. (C) The top panel shows the potentially opposite effect of two immune genes, *ADAR* and *DHX9*, on A- to-I RNA editing. The heatmap and correlation plots on the left display the co- existence of these two genes in 33 cancer types. The circle plot in the middle reveals the similar regions where these two proteins bind. The correlation plots on the right show the positive effect of *ADAR* and the negative effect of *DHX9* on the edited region of *PKR*. (E) The left histogram shows the number of innate immune genes associated with the edited regions of genes (x-axis). The heatmap and correlation plots display the relationships between the edited region of *PKR* and its own gene. The gene structure plot uncovers the possible mechanisms underlying the editing effects on genes. The histogram plot in the right panel shows the number of cancer types supporting the effects of this edited region on other interferon-stimulated genes (y-axis). The red color represents the positive effect, and the green color represents the negative effect respectively.

Particularly, these editing-associated genes cover interferon-stimulated genes (18) and RNA-binding proteins (19) (Figure 3B). They include the main editing enzyme of *ADAR* (20,21), famous RNA sensors such as *EIF2AK2* (*PKR*) (22) and *IFIH1* (*MDA5*) (23), and reported RNA-binding regulators of editing such as the DEAH box helicase 9 (*DHX9*) (24,25) and *ILF3* (26,27). A close look at these editing-gene associations and the intrinsic functions of genes discovers the interplay between A-to-I RNA editing and immune genes.

The well-studied editing-associated gene is *ADAR*. It is both an interferon-stimulated gene and an RNA-binding protein. This gene has more positive associations with the editing levels of informative regions (Figure 3B), describing the RNA regulatory function of one immune gene on A-to-I RNA editing. Moreover, other immune genes can also affect A-to-I RNA editing through the mediation of *ADAR*. For example, the *DHX9* protein, cooperating with *STAT1* to transcribe interferon-stimulated genes (28), interacts with *ADAR* to repress RNA editing (25). It is supported by the remarkable co-expression of these two genes in all the cancer types, the similar dsRNA substrates of these two proteins (24), and their effects on editing regions (Figure 3C).

On the other hand, A-to-I RNA editing sites prefer the location of non-coding regions, such as 3’-UTRs. There are 309 edited regions in 3’-UTRs significantly associated with the expression of innate immune genes. They may disturb the regulations of miRNAs and RNA-binding proteins on these immune genes (29,30). The edited region that has the relationships with the most immune genes locates in the famous RNA sensor of *PKR* (Figure 3D). It may cause the gain of 406 miRNA interactions, the loss of 339 miRNA interactions (29), and the regulatory intervention of RNA-binding proteins such as *DHX9* (19). Eventually, the editing events of the *PKR* gene lead to the downregulation of this gene in most cancer types, similar to that in Alzheimer’s disease and Parkinson’s disease (30). Further, due to the potential RNA-binding function of this protein (31), this edited region may affect additional immune genes, such as the downregulation of *TRIM25* and the upregulation of other interferon-stimulated genes in at least ten cancer types (Figure 3D). All the analyses introduce the potential effects of A-to-I RNA editing on immune genes.

From the above analyses, the immune genes and A-to-I RNA editing constitute complex mutual relationships. Importantly, the editing events in the 3’-UTR of *PKR* lead to the downregulation of this gene. Due to the crucial role of this gene in innate immunity (32), it reveals one potential mechanism of RNA editing as an inhibitor of *PKR*-related cancer immunity. Additional possible mechanism of A-to-I RNA editing in cancer immunity is studied further in the following parts.

### 4. RNA editing acts as a negative regulator of *MDA5*-activated interferon response

Interferons (IFNs) are pleiotropic cytokines critical for immune system regulation and immunotherapy response (5). The trigger factors of interferon-related signals include RIG-I-like receptors, such as *MDA5* (23). This gene recognizes dsRNA and then interacts with the adaptor *MAVS*, ultimately leading to the transcription of the genes encoding type I interferons (33,34). Due to the preferred dsRNA structure of A-to-I RNA editing (Figure 1E), it is necessary to explore the role of RNA editing in *MDA5*-activated interferon response for cancer.

The potential roles of interferons in cancers are first validated in more than ten cancer types with differential levels of interferon responses (35), as shown in Figure 4A and Figure S1A. The genes upregulated in response to interferon include *ADAR* and *PKR* (18). For example, more than 27 cancer types have remarkably positive associations between interferon signaling and the expressions of these two genes (Figure 4B, Figure S1B). The upregulated *ADAR* stimulated by interferons generates higher editing levels. It is supported by more positive *ADAR*-editing associations in TCGA samples and more down- edited sites in *ADAR* knockdown cell lines (Figure 4C, Figure S1C). The analyses show the effect of interferon on A-to-I RNA editing.

**Figure 4.**
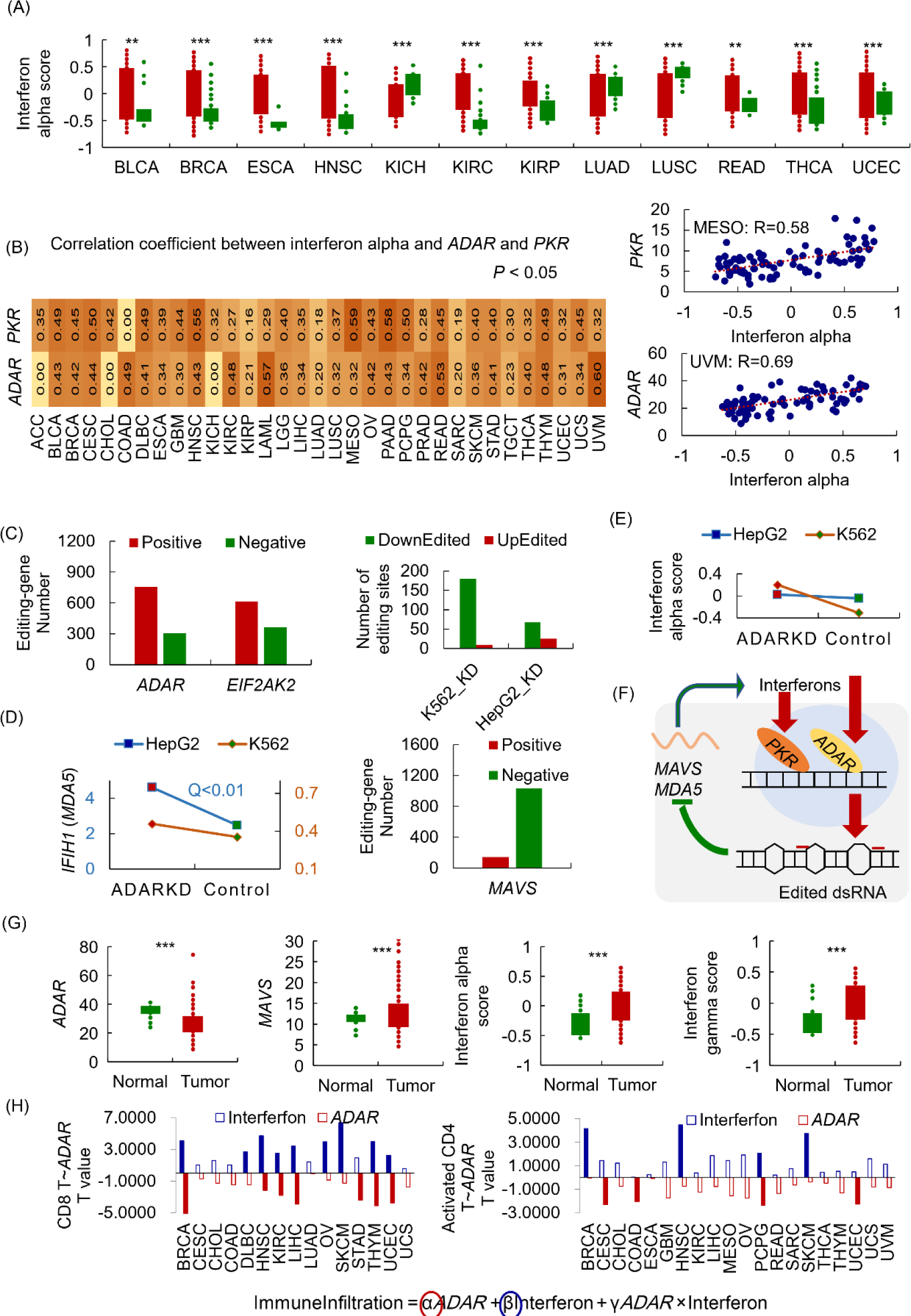
The effect of RNA editing on interferon response. (A) Differential alpha interferon score between tumor samples and controls for 12 cancer types. (B) The heatmap and correlation plots introduce two genes stimulated by alpha interferons in 33 cancer types. (C) The positive effect of *ADAR* on RNA editing levels in 33 cancer types (left) and *ADAR* knockdown cell lines (right). (D) The negative correlations between RNA editing and *MDA5* or *MAVS* in *ADAR* knockdown cell lines (left) and 33 cancer types (right). (E) The negative effect of RNA editing on alpha interferons in *ADAR* knockdown cell lines. (F) The diagram presents the hypothesis of the editing effect on interferon response. (G) In the KIRP cancer type, lower expression of *ADAR* upregulates *MAVS* expression and enhances interferons. (H) The contributions of *ADAR* and alpha interferon to the infiltration of CD8 T cells (left) and activated CD4 T cells (right).

Specifically, an up-regulation of *MDA5* is discovered in the low-editing cells (Figure 4D). In the cancer samples from TCGA, the downstream interacting gene of *MDA5*, *MAVS*, has the most significant associations with enriched editing regions. Of them, 87.67% presents negative correlations. This inverse relationship between A-to-I RNA editing and *MDA5*-related genes describes the inhibition roles of editing in *MDA5* activation (36). The editing repression on *MDA5* subsequently reduces the generation of interferons, supported by the enhanced interferon responses in *ADAR* knockdown cells (Figure 4E, Figure S1D). All the above analyses suggest RNA editing as a negative regulator of *MDA5*-activated interferon response (Figure 4F).

According to the analyses above, in vivo, a negative feedback loop exits to keep the homeostasis of interferons once activated. It covers the positive effect of interferons on A-to-I RNA editing and the negative regulation of *MDA5*-activated interferon response by RNA editing. One of the two-directional effects is superior in different cancer types, leading to significantly differential interferons in tumors (Figure 4A, Figure S1A). For example, in KIRP cancer, the lower expression of *ADAR* upregulates *MAVS* to enhance innate immunity by increasing interferon levels (Figure 4G).

With the interferon changes affected by A-to-I RNA editing, the fraction of immune cells alters in the tumor microenvironment. For example, *ADAR* negatively affects the infiltration of activated CD4 T cells and CD8 T cells in the 23 cancer types from TCGA, while interferon shows positive effects (Figure 4H). All the analyses reveal the negative regulation of RNA editing on interferon and interferon-related immune infiltration.

### 5. Low-editing patients benefit more from cancer immunotherapy

The above analyses uncover the potential function and mechanism of A-to-I RNA editing in tumors. One hypothesis suggests RNA editing as a pathogenic cancer biomarker, supported by the higher RNA editing levels in tumors compared to controls, tumors with higher stages, and poor survival patients (Figure 2). Another hypothesis reveals the inhibitor role of RNA editing in interferon-related cancer immunity with their interaction analysis results (Figure 4). The multivariate Cox regression analysis also validates this hypothesis with a favorable prognosis of interferons and a negative impact of *ADAR* on survival (Figure 5A). The potential roles of RNA editing in tumor promotion and cancer immunity guide a further study on immunotherapy. According to these two hypotheses, RNA editing may play a resistant role during cancer drug treatment.

**Figure 5.**
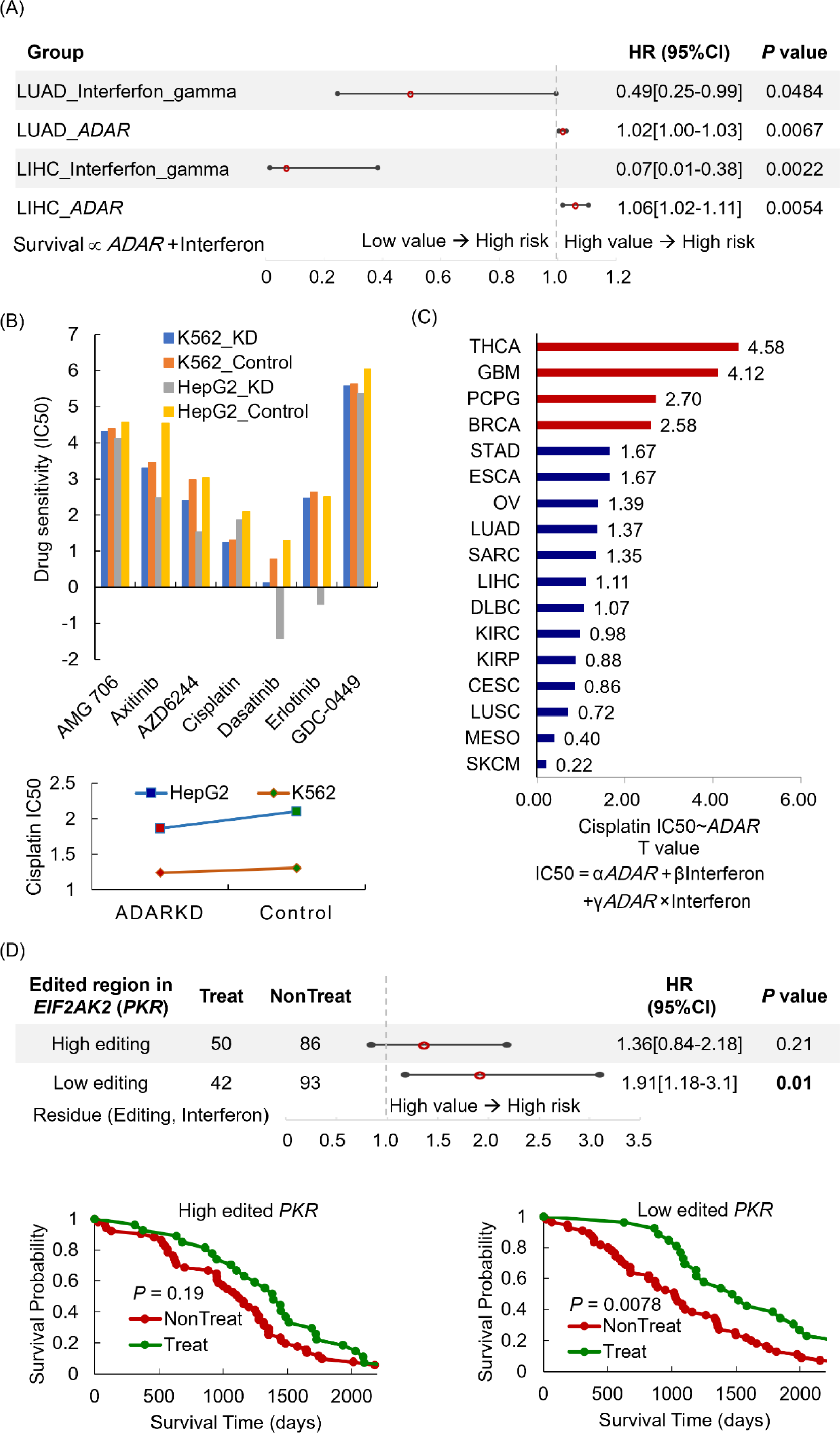
The effect of RNA editing on drug responses. (A) The multivariate Cox regression analysis results for *ADAR* and interferons in LUAD and LIHC cancer type. (B) The top histogram shows the sensitives of seven drugs between *ADAR* knockdown cell lines and wild types. The bottom panel displays a lower cisplatin concentration needed to inhibit tumor cells for *ADAR* knockdown cell lines. (C) The positive associations of IC50 with *ADAR* expressions by multiple linear regression analysis for patients from TCGA while interferons show negative associations. The red color represents significant associations (*P* < 0.05), and the blue color shows others. (D) The top panel shows the survival risk of patients with no cisplatin treatment in the low and high editing groups. The group is determined by the residues of a linear regression between editing levels of *PKR* and alpha interferons. The left bottom panel displays the KM survival analysis between patients treated with cisplatin and other patients, both of which have higher editing levels of *PKR*. The right bottom panel is KM survival analysis for the patients with lower editing levels of *PKR*. IC50, half maximal inhibitory concentration; KM, Kaplan-Meier.

The editing effect on drug responses is first validated by comparing the sensitivities of seven common cancer drugs between *ADAR* knockdown cell lines and wild types. A low concentration of drugs (IC50) is needed to inhibit tumor cells after *ADAR* knockdown (Figure 5B). These drugs include a chemotherapy medication of cisplatin for treating multiple cancers, such as ovarian cancer, cervical cancer, esophageal cancer, lung cancer, mesothelioma, and brain tumors. The positive associations between *ADAR* expression and required cisplatin concentration in these cancer types from TCGA (Figure 5C) describe the same story. The results reveal that these tumor patients with low *ADAR* expression benefit more from potential drugs.

The hypothesis still holds for specific enriched RNA editing regions, such as one in the famous immune gene of *PKR*. As shown in Figure 5D, patients with high editing levels show no significant improvement in survival after cisplatin treatment, while patients with low editing levels can benefit from cisplatin therapy. Thus, the downregulation of RNA editing may be a potential method to enhance drug responses by upregulating interferon-related cancer immune.

One last thing needed to be addressed is that there are possible side effects for the treatment design through the downregulation of RNA editing. The side effects may come from unfavorable roles of interferon in cancers (5,37). The persistent interferon signaling dampens immune responses and sustains resistance to DNA-damaging therapies (38). The interferon-mediated editing-related treatment thus may negatively affect drug responses. For example, in the head and neck cancer type, *ADAR* expression shows a favorable prognosis ability of survival, while interferon presents a negative effect (Figure S2). Another side effect of the editing-related treatment is the transition from epithelial to mesenchymal status (EMT). In all 21 cancer types from TCGA, *ADAR* may prevent the EMT process, while interferon displays promotion effects (Figure S3). One previous study also proposes the induction of EMT in *ADAR* knockdown cell lines (39). Thus, the downregulation of RNA editing to enhance interferon for tumor repression requires careful consideration for specific cancer and patient. Improper design of treatment may result in immune envision and tumor metastasis.

## Discussion

Here, we study the roles of A-to-I RNA editing during interferon pathways in tumors and its effect on cancer drug treatment from both view of *ADAR* and specific enriched editing regions (Figure 6). The regulators of A-to-I RNA editing include interferons stimulating the main editing enzyme and RNA-binding proteins such as *DHX9*. Both positive and negative effects on RNA editing exist in this process. Additionally, the downstream analysis of RNA editing reveals its inhibition of interferon through deactivating cellular dsRNA sensors such as *MDA5* (40) and *PKR*. These two analyses describe the complex interactions between A-to-I RNA editing and interferon. Due to the crucial roles of interferon in immune infiltration and cancer drug treatment (41,42), RNA editing negatively regulates the infiltration of CD4/ CD8 T cells and potentially decreases drug sensitivity through its effect on interferons.

**Figure 6.**
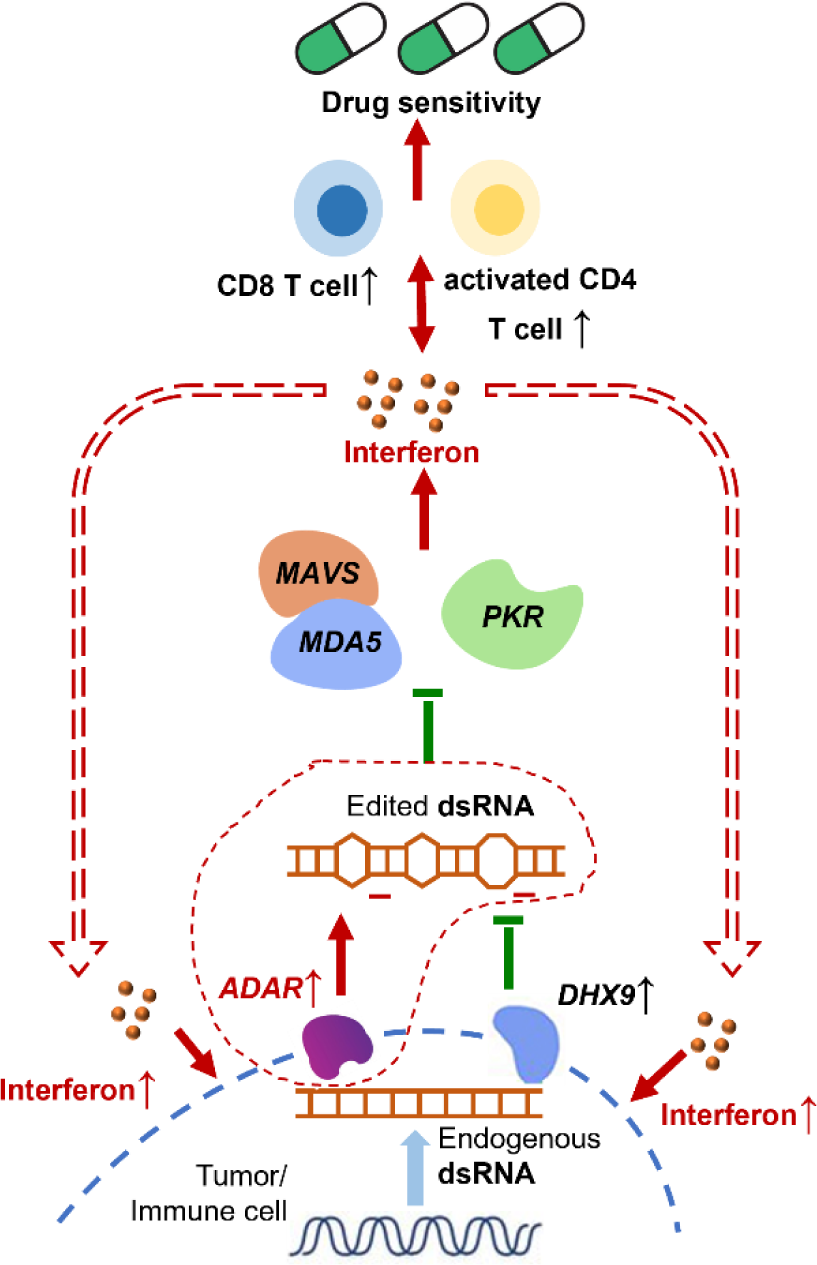
The roles of A-to-I RNA editing in interferon pathway and its effects on drug treatment design.

The downregulation of RNA editing is a potential way for cancer treatment. Its favorable aspects include the increase of interferon to enhance immune infiltration (Figure 4) and improve drug responses (Figure 5) (43). On the contrary, this treatment may also promote tumor progression through resistance to several cancer therapies (Figure S2) (44), epithelial-to-mesenchymal induction (Figure S3) (45,46), and CD8 T cell exhaustion (47). Therefore, RNA editing-related treatment requires careful design for specific cancer and patient in future clinics usage.

## Methods

### Detection of enriched editing region

There were three datasets involved in this study for the role of A-to-I RNA editing in cancer immunity. They included 11,052 patients across 33 cancer types from TCGA, kidney tissues from GEO (13), and *ADAR* knockdown cell lines (ENCSR164TLB, ENCSR667PLJ, ENCSR104OLN, and ENCSR305XWT) from ENCORE project. The A-to-I RNA editing sites and levels for each sample were either downloaded from CAeditome (29) or detected using a similar procedure. The preferred dsRNA genomic location for these editing events had close connections with the sensing regions of the proteins related to innate immunity (11). Then the analysis of RNA editing with cancer immunity started from the definition of enriched editing regions using the pipeline shown in Figure S4A. More samples in each cancer type provided more enriched editing regions (Figure S4B). Thus, this study used all the patients across 33 cancer types to define the enriched editing regions. Their distributions in diverse genomic locations and repeats were analyzed by ANNOVAR (48). Their dsRNA structures were validated by the blast alignment (49) between the enriched editing regions and the surrounding sequence fragments (< 1e4).

### Analysis of tumor-specific enriched editing region

The tumor-specific enriched editing regions can reveal the trend of A-to-I RNA editing in tumor samples, severe cancer patients, and patients with poor survival. For this goal, the editing levels between tumors and adjacent normal controls were compared with the t-test method (*P* < 0.05, informative number >= 20) to identify the enriched editing regions associated with tumor occurrence. The editing levels correlated with pathologic stages by both ANOVA (*P* < 0.05) and Pearson method (*P* < 0.05, informative number >= 20) for the edited regions related to tumor progression. The survival time for patients with low and high editing levels was differentiated with both the log-rank test (*P* < 0.05) and Cox analysis (*P* < 0.05) to recognize the edited regions associated with cancer survival.

### Correlation of enriched editing regions and immune genes

The analysis of cancer immunity started from the correlations of enriched editing regions with immune genes. Then quantitative trait locus (QTL, FDR < 0.05) and Pearson (*P* < 0.05) tests were used sequentially to identify the enriched editing regions associated with gene expressions. Common confounders, such as gender, age, and disease stages, were combined as the covariates for QTL analysis to improve the identification sensitivity. The immune genes got from InnateDB (15). Their involved biological functions were provided with Enrichr (50).

### Correlation of enriched editing regions and immune pathways

The immune pathways involved in this study included interferon alpha response, interferon gamma response, and epithelial-mesenchymal transition from MSigDB (35). They were evaluated by the GSVA method (51). The scores of interferon responses correlated with the enriched editing regions by the Pearson method (*P* < 0.05). The scores of epithelial-mesenchymal transition linked with the enriched editing regions by multiple joint regression analysis (*P* < 0.05). A different method used here was due to the potential effect of mutual interactions between editing and interferon (18) on other immune pathways (39,45).

### Correlation of enriched editing regions and immune infiltration

For each patient, the fractions of immune cells were estimated by CIBERSORT (52). They correlated with the enriched editing regions by multiple joint regression analysis (*P* < 0.05) due to the potential effect of mutual interactions between editing and interferon (18) on immune cells (41,53).

### Correlation of enriched editing regions and cancer immunotherapy

The sensitivities of tumor drugs for each sample were estimated with the index of half maximal inhibitory concentration (IC50) by pRRophetic package (54). The higher IC50 value described a lower drug sensitivity in general. It correlated with the enriched editing regions by multiple joint regression analysis (*P* < 0.05). Especially for cisplatin with sufficient information from TCGA, the survival probability was compared between patients treated with cisplatin and patients untreated with cisplatin in both high and low editing groups. The differential treatment effectiveness provided additional evidence for the editing effect on drug responses.

## Author Contributions

Conceptualization, S.W.; methodology, S.W.; formal analysis, S.W.; investigation, S.W., J.W.; visualization, X.Q., Z.L., P.K.; data curation, S.W., M.Y.; writing—original draft preparation, S.W.; writing—review and editing, J.W., X.Z.; supervision, X.Z. and L.H.; funding acquisition, S.W. and L.H. All authors have read and agreed to the published version of the manuscript.

## Informed Consent Statement

The data was obtained from public resources of The Cancer Genome Atlas (TCGA), Gene Expression Omnibus (GEO), and ENCORE project.

## Data Availability Statement

The data presented in this study are available in Supplementary Material here and https://relab.xidian.edu.cn/ImmuneEditome/.

## Acknowledgement

The results <PUBLISHED or shown> here are in whole or part based upon data generated by the TCGA Research Network: https://www.cancer.gov/tcga. This research was funded by the National Natural Science Foundation of China (Grant No. 62002270), the Fundamental Research Funds for the Central Universities, the National Natural Science Foundation of China (Grant No. 82227802), the Natural Science Foundation of Shaanxi Province of China (Grant No. 2020JQ-332), the China Postdoctoral Science Foundation (Grant No. 2018M643583), and the National Key R&D Program of China (Grant No. 2017YFA0205202) and partially funded by the National Natural Science Foundation of China (Grant No. 61672422).

