## Supplementary Figure 1-4 for "The roles of RNA editing in cancer immunity through interacting with interferon"

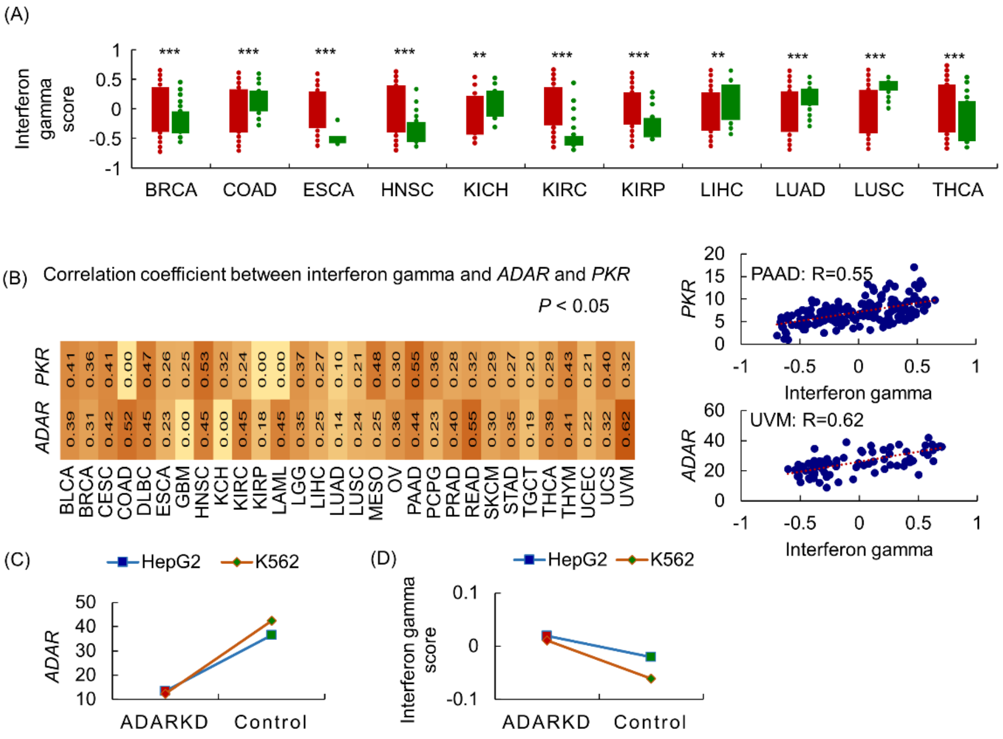


Figure S1. The effect of RNA editing on gamma interferon response. (A) Differential gamma interferon score between tumor samples and controls for 11 cancer types. (B) The heatmap and correlation plots introduce two genes stimulated by gamma interferons in 30 cancer types. (C) The validation of *ADAR* expressions in *ADAR* knockdown cell lines compared to controls. (D) The negative effect of RNA editing on gamma interferons in *ADAR* knockdown cell lines.

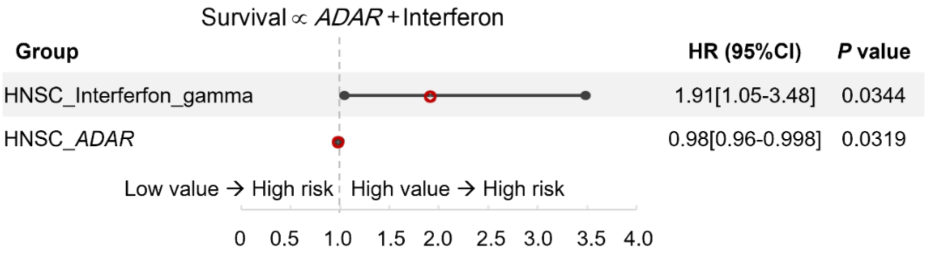


Figure S2. The multivariate Cox regression analysis results for *ADAR* and interferons in HNSC cancer type.

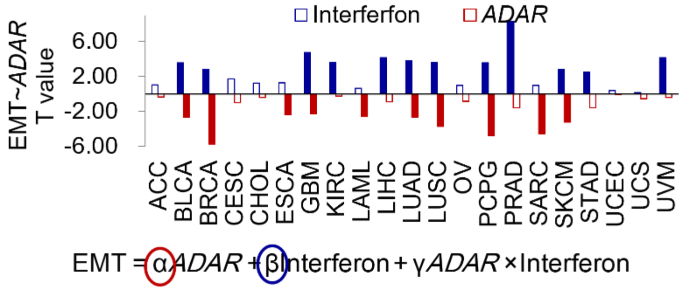


Figure S3. The contributions of alpha interferons and *ADAR* on EMT. EMT, epithelial-mesenchymal transition.

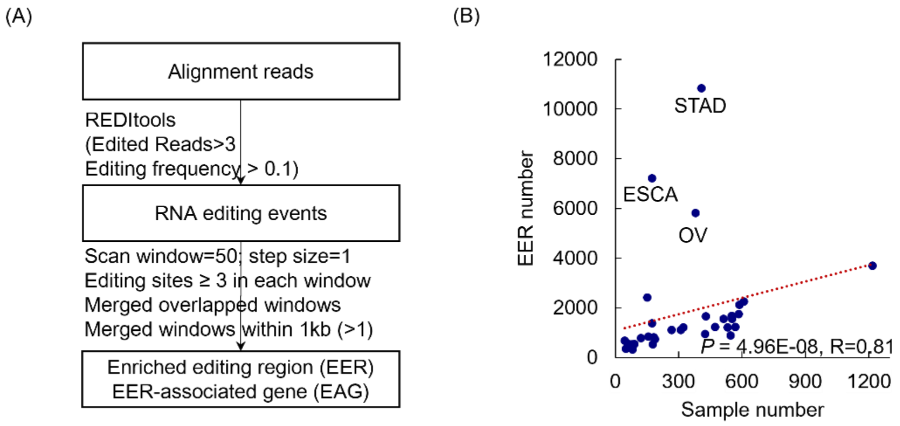


Figure S4. The definition of enriched editing regions. (A) The pipeline to identify enriched editing regions. (B) The correlations between the number of enriched editing regions and the number of samples for each cancer type.
